# Predicting molecular recognition features in protein sequences with MoRFchibi 2.0

**DOI:** 10.1101/2025.01.31.635962

**Authors:** Nawar Malhis, Jörg Gsponer

## Abstract

Molecular Recognition Features (MoRFs) are segments within disordered protein regions (IDRs) that undergo a disorder-to-order transition upon binding to their partners. Identifying MoRFs remains a significant challenge. This paper introduces MoRFchibi 2.0, a specialized prediction tool designed to identify the locations of MoRFs within protein sequences. Our results show that MoRFchibi 2.0 outperforms all existing MoRF and general predictors of protein-binding sites within IDRs, including top-performing models from CAID rounds 1, 2, and 3. Remarkably, MoRFchibi 2.0 surpasses predictors that utilize AlphaFold data and state-of-the-art protein language models, achieving superior ROC and Precision-Recall curves and higher success rates. MoRFchibi 2.0 generates output scores using an ensemble of logistic regression convolutional neural network models, followed by a reverse Bayes Rule to adjust for priors in the training data. These scores reflect MoRF probabilities normalized for the priors in the training data, making them individually interpretable and compatible with other tools utilizing the same scoring framework.

**Availability:** An online server: https://mc2.msl.ubc.ca/index.xhtml and code: https://github.com/NawarMalhis/MC2.git.

## 1 Introduction

Intrinsically disordered regions (IDRs) in proteins are segments that lack a defined tertiary structure under physiological conditions ^[1]^. Despite this, IDRs are functionally significant and often play key roles in cellular processes by interacting with other proteins, DNA, RNA, or small molecules ^[2-5]^. Their structural flexibility and dynamic nature allow IDRs to adapt and mold themselves around their interaction partners ^[6, 7]^.

As of the latest release of the DisProt database ^[8]^ in June 2024, 641 out of 3,022 proteins annotated with IDRs include one or more experimentally validated binding segments. However, this is a small number, especially considering that approximately 32% of the human proteome consists of proteins with more than 30% disordered residues ^[9]^. Consequently, the computational identification of binding regions within IDRs has emerged as a significant challenge. Of particular interest are regions within IDRs that bind to other proteins, and numerous computational tools have been developed to aid in identifying these protein-binding segments ^[10-46]^.

A subfamily of protein-binding segments in IDRs is known as molecular recognition features (MoRFs) ^[47-49]^. These segments transition from a disordered to an ordered state upon binding to their interaction partners. Typically, MoRFs are short, though they can sometimes span 50 or more residues. Many MoRF-mediated interactions are characterized by low affinity but high specificity, making them well-suited for reversible regulatory interactions in signaling pathways. The functional significance of MoRFs, along with the challenges posed by their experimental identification and characterization, has led to the development of several computational tools designed to predict the locations of MoRFs in protein sequences. Notable examples include Retro-MoRFs ^[13]^, MoRFpred ^[18]^, fMoRFpred ^[23]^, MoRFchibi_light ^[25]^, MoRFchibi_web ^[21]^, OPAL ^[31]^, MoRFPred-plus ^[29]^, and MoRFcnn ^[44]^. Since MoRFs represent only a small fraction of all residues in protein sequences, i.e., low “priors” probabilities, we require strong evidence, i.e., high-quality predictors, to achieve acceptable posterior probabilities ^[50]^, i.e., precision.

MoRFchibi predictors, light and web, are among the most accurate MoRF predictors available today. They are ranked among the leading IDR-protein binding predictors in the first two rounds of the CAID competition ^[51, 52]^ and are available among the CAID Portal set of predictors ^[53]^. MoRFchibi light predictions are included in the DescribePROT database ^[54]^, discussed in several review articles ^[55-59]^, and utilized in various applications ^[60-63]^. However, the accuracy of predictions depends heavily on the quality and size of the training datasets, and MoRFchibi predictors were developed using training data collected in 2008 ^[18]^. Since then, several databases have been created that contain curated and annotated IDR protein-binding segments, including MoRF segments. The availability of more extensive and accurate training data enabled us to redesign MoRFchibi and increase the quality of its prediction. Here, we introduce MoRFchibi 2.0, a redesigned MoRF predictor that encompasses an ensemble of four logistic regression convolutional neural network (LR-CNN) models. We also provide newly assembled datasets with MoRF annotations for the community to optimize and test other models.

## 2 Methods

### 2.1 Datasets

We assembled a dataset of protein sequences annotated with disordered regions that undergo disorder-to-order transitions upon binding to other proteins, drawing from four manually curated databases: DisProt, DIBS ^[64]^, MFIB ^[65]^, and IDEAL ^[66]^. Two key factors that impact the quality of MoRF annotations are the availability of evidence supporting the disorder of a sequence segment in isolation and proof of folding induced by binding to a protein target. The extent to which this critical information is covered varies across the four databases.

DisProt, a database for disordered protein sequences, provides disorder annotations and details on IDR properties, including protein and nucleotide binding, as well as disorder-to-order (DtO) transitions. DIBS and IDEAL are databases that focus on interactions between binding segments in IDRs and their ordered protein partners. In these databases, the disordered state of binding segments is supported either by experimental data or inferred from homology. While all interacting IDR segments in IDEAL undergo a disorder-to-order transition upon binding, a small fraction of these segments in DIBS do not fold upon binding. The MFIB database, on the other hand, includes IDRs that bind to other IDRs and fold. The majority of interactions in MFIB are backed by experimental evidence, with some based on homology.

Using the latest DisProt release (24_06), we annotated all disordered protein-binding residues that overlap with DtO transitions as MoRF residues. Since DtO annotations are likely incomplete, we masked out annotated protein-binding residues not associated with DtO, meaning we considered them neither MoRFs nor non-MoRFs. We included IDEAL and DIBS binding segments that were validated using experimental data and restricted the DIBS segments to those associated with DtO annotations in DisProt. Additionally, we incorporated all MFIB binding annotations. This process resulted in 899 high-quality MoRF segments (HQ MoRFs) across 769 sequences, encompassing 46,694 residues. Alongside these HQ MoRF annotations, we included low-quality MoRF annotations (LQ MoRFs) for training. These LQ MoRFs are binding segments from IDEAL and DIBS, supported by evidence from homology and DIBS binding segments not corroborated by DisProt DtO annotations. This approach yielded 1,217 MoRF segments (HQ and LQ) across 1,030 sequences, totaling 52,560 residues. Thus, 769 sequences are annotated by HQ MoRFs and possibly LQ MoRFs, and the remaining 261 sequences are annotated by LQ MoRFs only.

A high degree of redundancies in the testing data can lead to biased evaluations. In the training data, it leads the optimization process to over-weight these redundant sequences at the expense of others. Thus, we used CD-Hit ^[67]^ to cluster these sequences at an 80% identity. To avoid unnecessarily losing some training MoRF annotations, within each cluster, MoRF annotations for sequences with >95% identity to cluster centers or >92% identity and equal length to cluster centers are transferred to cluster centers as LQ annotations, i.e., to be used for training only. Then, we only used cluster centers, resulting in 907 sequences with 1,081 MoRFs, totaling a combined 46,504 HQ and LQ MoRF residues. This included 674 sequences with 790 HQ MoRF segments, totaling 40,657 MoRF residues.

As sequences sourced from the DisProt database are likely to be more completely annotated than those from IDEAL, DIBS, and MFIB, the non-MoRF residues in DisProt sequences are less likely to include false negatives. Thus, we masked out non-MoRF residues in sequences not sourced from DisProt.

We divided the total 907 MoRF sequences into five subsets such that sequences in each subset are less than 40% identity to those in other subsets. For that, we clustered the sequences at 40% identity using CD-Hit. We sorted these clusters in descending order based on the total number of HQ MoRF residues in each cluster. Then, starting with five empty subsets, we iteratively moved clusters to these subsets, such that at each iteration, we moved the cluster with the highest number of HQ MoRF residues to the subset with the least total number of HQ MoRF residues.

We used four subsets containing both LQ & HQ MoRF annotations to train four cross-validated LR-CNN models and the remaining fifth subset for testing. We further filtered out sequences in the test dataset with an identity higher than 30% to those in the four training datasets, resulting in a test dataset with 143 sequences (Test-143). Although sequences in the test dataset are less than 30% identical to those used in the training, short test regions can still have local homology to training sequences. Thus, we used HAM ^[68]^ to identify and mask out regions with 10 or more residues in the test sequences that are homologs (with an identity of greater than 80%) to the training sequences. HAM identified three regions (totaling 33 residues) in the testing dataset as homologs of the training data, one of which is a homolog of a masked training region of 11 residues. Therefore, HAM only masked the remaining 22 residues. Thus, the test set (Test-143) contains 143 sequences with 148 MoRFs, totaling 7,757 MoRF residues. For completeness, we also created Test-143 (unmasked), for which all previously masked residues, except those masked by HAM, were unmasked. In addition, we created the Test-128 (AF2) subset, which is limited to Test-143 sequences with AlphaFold2 structures in the AlphaFoldDB database ^[69]^. We also created the Test-106 (SR) subset, which only contains Test-143 sequences with both MoRF and non-MoRF residues. Specifically, we excluded all sequences from Test-143 that do not have at least five residues for each class, i.e., all sequences must have five or more MoRF residues and five or more non-MoRF residues. Moreover, as SR is not sensitive to under-annotated sequences and to avoid significantly reducing the test data size, we unmasked non-MoRF sequences outside DisProt. **Table 1** describes the details of all four training datasets, the Test-143 dataset, and the three variations.

**Table 1:** Training and testing datasets. Four subsets, cv1, cv2, cv3, and cv4, utilize LQ annotations for training and HQ annotations for validation. And the fifth only used HQ annotations for testing.

|  | MoRFs quality | Sequences with MoRFs/Total | MoRFs | Non-MoRF Residues | MoRF Residues | Masked Residues |
| --- | --- | --- | --- | --- | --- | --- |
| **Training (CV1)** | LQ & HQ | 141/141 | 172 | 28,812 | 8,902 | 47,440 |
| **Training (CV2)** | LQ & HQ | 182/182 | 208 | 57,212 | 9,267 | 43,553 |
| **Training (CV3)** | LQ & HQ | 192/192 | 232 | 51,231 | 9,347 | 45,729 |
| **Training (CV4)** | LQ & HQ | 198/198 | 237 | 52,908 | 9,532 | 58,255 |
| **Test-143** | HQ | 126/143 | 148 | 46,744 | 7,757 | 20,037 |
| **Test-143 (unmasked)** | HQ | 126/143 | 148 | 66,759 | 7,757 | 22 |
| **Test-128 (AF2)** | HQ | 113/128 | 132 | 33,923 | 7,246 | 16,449 |
| **Test-106 (SR)** | HQ | 106/106 | 128 | 49,676 | 5,823 | 1,163 |

### 2.2 Input Features

We used the seven features introduced in ^[45]^ to encode amino acid sequences. The first two features estimate the energy associated with intrachain interactions of each residue by using the previously derived 20 × 20 energy predictor matrix ^[70]^. Specifically, the first interaction energy feature is the sum of the pairwise energies between the residue at k and all residues within a window of w1 = 10 on either side of k. The second interaction energy feature is the sum of this energy for all residue pairs between w1 and w2 = 25 residues on either side of k. The additional five features were obtained from the **F5** amino acid enrichment matrix. **F5** is a 20 × 5 matrix encoding the enrichment of each amino acid within IDRs, linker regions, IDR nucleic binding sites, IDR protein binding sites, and folded domains, as described in ^[45]^.

### 2.3 Model selection and optimization

We used the ensemble window-in/window-out structure utilizing four LR-CNN models introduced in ^[45]^. An input sliding window size ten is padded by the 250 residues on each side, totaling 510 window-in residues. If not enough residues are available for padding in the input sequence, blank residues are used in their place. A blank residue is a hypothetical residue with all its features are zeros. We trained four LR-CNN models, each on three CV subsets, and validated them on the fourth (see **Table 1** for details). Each model comprises three convolutional layers with a kernel size of seven, separated by two average pooling layers with a kernel size of seven, followed by two fully connected (FC) layers with the output size of the first FC layer set to 150. All the above hyperparameters were selected using a grid search to minimize the loss against the validation sets. We used a binary cross-entropy with a sigmoid (BCEWithLogitsLoss) as the training loss and applied a sigmoid function to the evaluation output of each LR-CNN model. Thus, each LR-CNN is a logistic regression model. We applied reverse Bayes to each model output to factor out the training priors, i.e., the imbalances in the percentages of MoRF residues between the training data subsets. Then, we averaged the four scores and used a smoothing window of equal weights size 5 to smooth the final output score. We trained 15 instances for each of the four LR-CNN models, each minimizing the validation loss. To avoid overfitting the validation subsets, we then selected the instance with the third-best validation AUC.

Finally, we need to highlight two crucial design issues: first, we are using a large padding size, 250 residues on each side, to provide the model with a larger receptive field, thus enabling a larger environment to be used in the evaluation whenever needed. Second, we employed an ensemble of four LR-CNNs trained on overlapping datasets to increase prediction consistency and mitigate potential biases that could arise from the limitations of our training data size.

### 2.4 Evaluation measures

We calculated Receiver Operating Characteristic (**ROC)** curves to evaluate the performance of tools independent of the binary threshold cut-off values. ROC curves chart the relationship between the percentage of the positive class passing any score (true-positive rate, TPR) and the percentage of the negative class passing that score (false-positive rate, FPR). The formal definitions of TPR and FPR are:

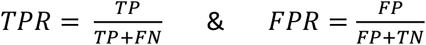

TP, FP, TN, and FN represent the counts of True Positives, False Positives, True Negatives, and False Negatives, respectively. We use the area under the ROC curve (AUC) values to evaluate the overall performance of a predictor in a single value. The diagonal line is the ROC curve for a random (naïve) classifier. The AUC for a random classifier is 0.5, and the better the classifier is, the higher its AUC value will be. Since ROC curves are insensitive to dataset imbalance, so are AUC values. For a randomly selected positive class instance and a randomly selected negative class instance, the AUC is the probability of ranking the positive instance higher than the negative one ^[71]^. We used the Precision-Recall curves to chart the relationship between precision and recall where:

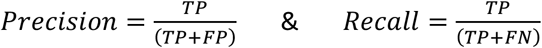

We included the average precision score (APS) ^[52]^ for each MoRF predictor in the legends of the PR curves. We also used the success rate (SR) to evaluate our predictor. SR measures the percentage of protein sequences with the average score of the positive class residues in the sequence (MoRF residues) higher than the average score of the negative class (non-MoRF residues). SR scores are essential as they are insensitive to under-annotation in the data and factor out the contribution of protein-level features that adjust the scores of residues for entire proteins based on the overall protein activity. Using protein-level features, such as protein-protein interaction data, can boost the AUC value of predictors by upscoring entire sequences of hub proteins. However, the outputs of predictors relying on protein-level features are less precise in identifying the location of MoRF residues within each sequence. Predictors with higher SR scores are more suitable for determining the position of the MoRF(s) in a protein sequence.

## 3. Results

### 3.1 Prediction assessment

We compared MoRFchibi 2.0 (MC2) predictions with those available for MoRF-specific and more general predictors of protein-binding regions in intrinsically disordered regions (IDRs). Specifically, we used the complete Test-143 set to compare MC2 predictions with those of the original MoRFchibi web and light methods, as well as those MoRF predictors that participated in the CAID1 or CAID2 competitions, such as fMoRFpred and OPAL. In addition, we compared predictions with those of thirteen methods that identify protein-binding segments in IDRs, independent of whether the segments undergo DtO transitions upon binding. **Figure 1** and **Table 2** reveal that MoRFchibi 2.0 outperforms all 17 tested predictors in terms of AUC and APS. In the Test-143 set, we masked all low-quality MoRF sites to reduce the likelihood of false negatives. To align with test datasets commonly used in the field and assess the impact of this procedure, we unmasked the LQ MoRF sites, as well as non-MoRF residues outside DisProt sequences, and created an additional test dataset. Results for this Test-143 (unmasked) dataset show a slight decrease in the AUC and APS of most predictors, including all specialized MoRF predictors, but no substantial change in the ranking in AUC performance. A recent study has shown that AlphaFold2 often assigns higher pLDDT scores to residues in MoRFs, residues that undergo DtO upon binding, and proposed that AlphaFold2 can identify MoRFs with high precision ^[42, 72]^. Therefore, we included two additional predictors (AlphaFold-Bind and IPA-AF2_protein) that rely on AlphaFold-2 structures in their predictions, specifically AlphaFold-2’s pLDDT scores ^[42, 45]^. Thus, we excluded sequences in the Test-143 dataset that do not have a structure available in the AlphaFoldDB (see **Table 1**). Results in **Table 2** show that MoRFchibi 2.0 continues to outperform all predictors, including those that use AlphaFold-2 scores. Finally, we compared predictors using success rates. To do so, we used the test dataset, Test-106 (SR), which only contains sequences with both MoRF and non-MoRF residues. Results in **Figure 1** and **Table 2** reveal that MoRFchibi 2.0 provides predictions for the Test-106 (SR) subset, with the highest AUC and SR values for all predictors tested. Note that while DisoFLAG-PR (dotted Brown line) has an AUC for this subset that almost equals that of MoRFchibi 2.0, the DisoFLAG-PR AUC value is the result of a better performance in identifying non-MoRF residues (top right corner) when compared to MoRFchibi 2.0, which is better in identifying MoRF residues (lower left corner of the ROC curve).

**Table 2:** Comparison of prediction performances. The AUC values for MoRFchibi 2.0 and other predictors using the complete test dataset, Test-143, the complete unmasked dataset, Test-143(unmask), the subset of the test dataset with AlphaFold structures, Test-128(AF2), and the subset of sequences with at least 5 MoRF and 5 non-MoRF residues, Test-106(SR). We also used the Test-106 (SR) subset to compare the success rates of these predictors. MoRF predictors are in bold, and the highest AUC and Success Rate values are underlined.

| Measure | AUC | AUC | AUC | AUC | Success Rate |
| --- | --- | --- | --- | --- | --- |
| Test Data | Complete | Unmasked | Subset (AF2) | Subset (SR) | Subset (SR) |
| **MoRFchibi 2.0** | <u>**0.813**</u> | <u>**0.803**</u> | <u>**0.806**</u> | <u>**0.745**</u> | <u>**0.736**</u> |
| **OPAL** | **0.759** | **0.752** | **0.744** | **0.683** | **0.660** |
| **MoRFchibi_web** | **0.703** | **0.702** | **0.710** | **0.653** | **0.717** |
| **MoRFchibi_light** | **0.698** | **0.694** | **0.700** | **0.636** | **0.689** |
| **fMoRFpred** | **0.597** | **0.591** | **0.597** | **0.579** | **0.670** |
| IPA_all | 0.750 | 0.722 | 0.729 | 0.703 | 0.708 |
| IPA_protein | 0.741 | 0.716 | 0.720 | 0.705 | 0.717 |
| DisoFLAG-PB | 0.727 | 0.715 | 0.726 | 0.741 | 0.726 |
| ProBiPred_protein | 0.675 | 0.676 | 0.673 | 0.627 | 0.575 |
| AIUPred-Binding | 0.629 | 0.610 | 0.623 | 0.645 | 0.632 |
| ENSHROUD_all | 0.628 | 0.606 | 0.645 | 0.644 | 0.632 |
| ENSHROUD_protein | 0.601 | 0.577 | 0.614 | 0.617 | 0.642 |
| DeepDISObind_all | 0.608 | 0.591 | 0.591 | 0.588 | 0.557 |
| DeepDISObind_protein | 0.598 | 0.582 | 0.581 | 0.581 | 0.632 |
| bindEmbed21IDR_idrGeneral | 0.609 | 0.618 | 0.604 | 0.576 | 0.481 |
| bindEmbed21IDR_rawGeneral | 0.601 | 0.611 | 0.595 | 0.569 | 0.377 |
| DisoRDPbind | 0.580 | 0.563 | 0.578 | 0.609 | 0.670 |
| ANCHOR2 | 0.575 | 0.556 | 0.566 | 0.600 | 0.613 |
| AlphaFold_binding |  |  | 0.657 |  |  |
| IPA-AF2_protein |  |  | 0.712 |  |  |

**Figure 1:**
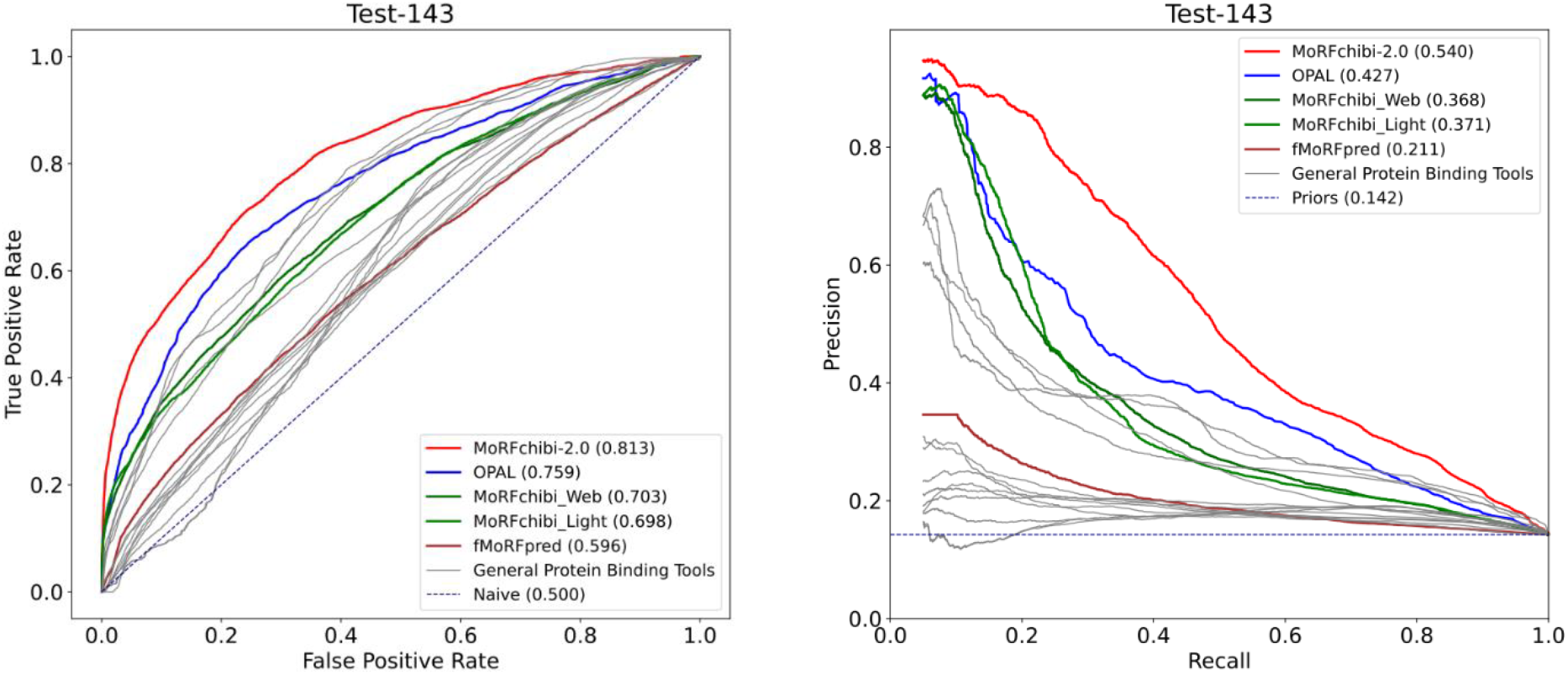
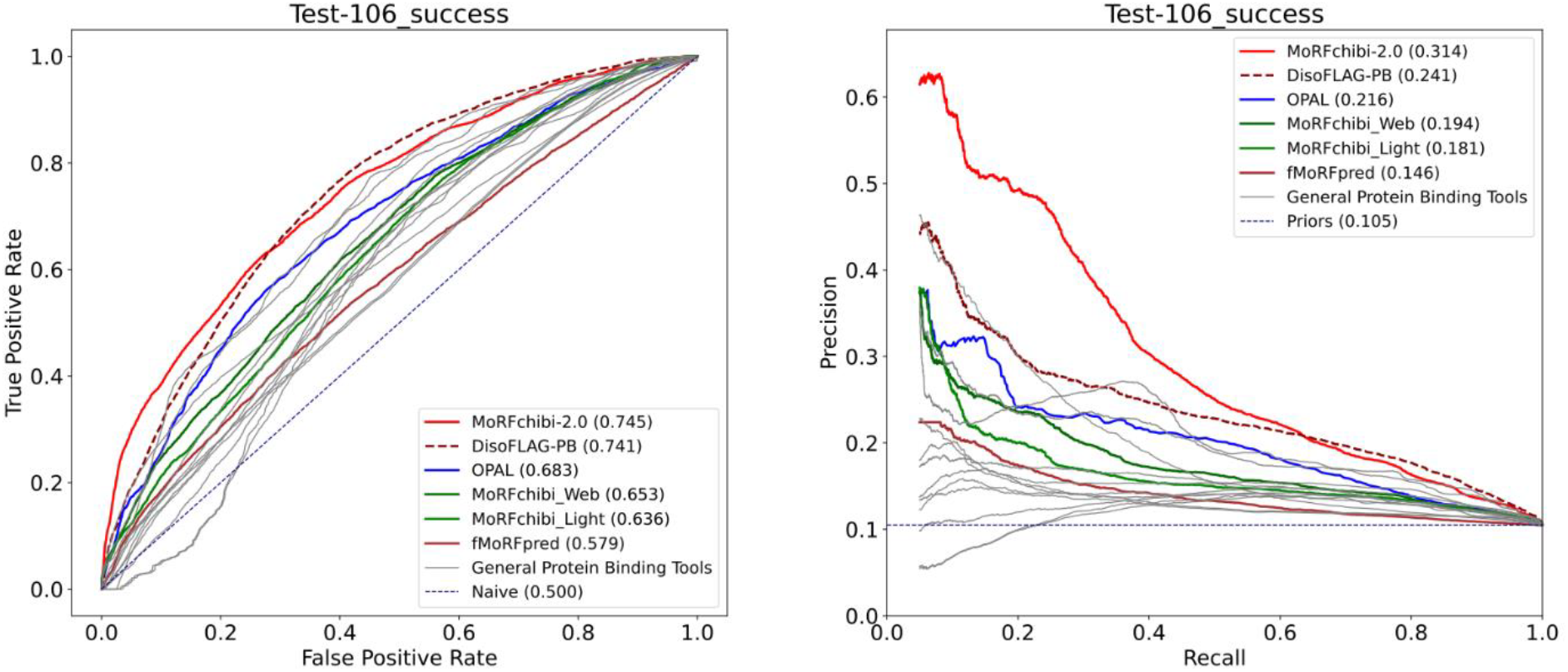
ROC and PR curves generated with predictions of MoRFchibi 2.0 and various methods. Top) MoRFchibi 2.0 performance (left: ROC and right: Precision-Recall) against the Test-143 dataset compared to a set of MoRF predictors: OPAL, MoRFchibi_web, MoRFchibi_light, and fMoRFpred, and thirteen general predictors of protein-binding segments in IDR (in gray lines). **Bottom)** the performance of these predictors against the SR subset, where the general binding predictor DisoFLAG-PB is in the doted Broun line to highlight MoRFchibi 2.0 higher performance at the most critical lower left corner of the ROC curve and the left of the PR curve. In the PR curves (Top and Bottom), to improve the clarity of the figures, we excluded points with recall values less than 0.05. The dotted gray lines represent the performance of a naïve predictor.

### 3.2 Case studies

To demonstrate MC2’s output and help understand its predictions, we present prediction examples for six of the Test-143 proteins in **Figure 2**. For the example in **Figure 2A**, MC2 successfully identifies the 157 residues annotated by DisProt as a MoRF. It is clear from the ROC and PR curves in **Figure 1** that such “perfect” predictions are not always possible, and there are scenarios where MoRFchibi 2.0 is not able to identify MoRF sites, as illustrated in **Figure 2B**, where MC2 only identifies residues 20-30 that are annotated in the DIBS database as MoRF and fails to identify the extended site 142-485 annotated as MoRF in DisProt. Not all cases are clear-cut; in **Figure 2C**, MC2 only identifies half of the MoRF annotated in DisProt (the residues 1-105 case). However, the paper cited by DisProt states that ‘the minimal AID construct σ54(16-41) is sufficient to form the activator complex’ ^[73]^, i.e., indicating that the first half of this site is more important than the second, notably consistent with MC2’s score distribution for this site. A similar case is shown in **Figure 2D**. While the segment 412-490 is annotated as MoRF in DisProt, a structure of the interacting MoRF is only available for the segment 465-490, where the MC2 score is the highest. Perhaps MoRF annotations should not be binary. In **Figure 2E**, MC2 scores suggest a MoRF between residues 378 and 439. However, DisProt only annotated residues 378-393 and IDEAL residues 435-439 as MoRFs, leaving the region 394-434 as an over-prediction by MC2 or an under-annotated region. Interestingly, MC2 seems to predict regions that undergo DtO upon binding to proteins and other molecules, which can be considered over-predictions. **Figure 3F** shows a case where MC2 scores identify the site around the 41-residue-long segment 294-334 as a MoRF. However, this 41-residue site, which binds to a protein partner, constitutes a zinc finger not stabilized by binding to the protein partner but by zinc ions ^[74]^.

**Figure 2:**
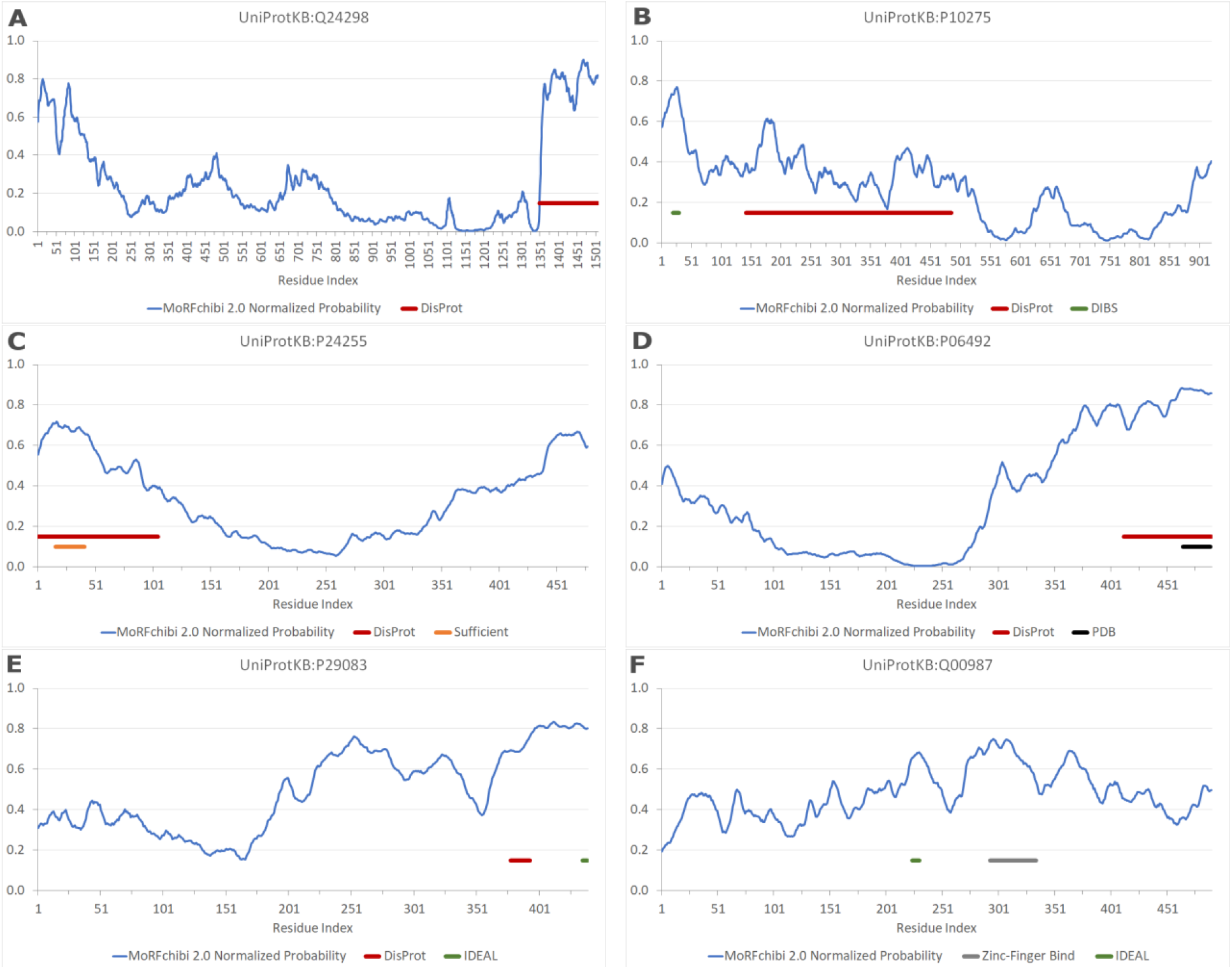
MoRFchibi 2.0 Normalized Probability scores in blue compared to the DisProt MoRF annotations in red, IDEAL and DIBS annotations in green, and those derived from PDB structures are in black. In (C), we also included a subset of DisProt annotations provided by the source publication, which were deemed sufficient to form the protein complex in Orange.

## 4 Discussion

We introduce MoRFchibi 2.0 (MC2), a specialized prediction tool designed to identify the locations of Molecular Recognition Features (MoRFs) in protein sequences. MoRFs constitute a subset of all binding regions within IDRs, those undergoing a disorder-to-order transition upon binding to protein partners. Our results demonstrate that MC2 outperforms existing MoRF predictors and predictors that identify protein-binding segments more broadly within IDRs. The latter group of predictors includes those that were ranked top in the community-driven critical assessments of protein intrinsic disorder predictions (CAID) rounds 1, 2, and 3 ^[75]^. Importantly, MC2 prediction performance surpasses even those methods that leverage AlphaFold output and use advanced protein language models, delivering superior ROC and Precision-Recall curves and higher success rates. This result is notable, given recent studies showing that segments within IDRs that undergo DtO transitions often have higher pLDDT scores than the surrounding segments or IDRs that do not generally have MoRFs ^[72]^. Regardless, neither MoRFchibi 2.0 predictions nor MoRF annotations are perfect. We provide examples that demonstrate MC2 predictions in various scenarios, including accurate predictions, under-predictions, over-predictions, and predictions for regions with ambiguous or incomplete annotations. Most importantly, MC2, relying solely on amino acid sequences, predicts regions with a low disorder ‘signature’ that bind to proteins and are likely to fold upon binding as MoRFs. However, we highlighted in **Figure 2F** that protein binding and protein folding can be influenced by environmental factors, such as ions or post-translational modifications, which can trigger a protein region to fold independently and mislead MC2 predictions.

MoRFchibi 2.0 generates output scores using an ensemble of four LR-CNN models followed by a reverse Bayes Rule. Consequently, these four scores, along with their average, represent probabilities normalized for the priors in the training data. As a result, MC2 scores are individually interpretable, enabling direct comparison and integration with other scores following the same output framework, such as IPA ^[45]^. We provide an HTML server and downloadable code for the broader community to use this new method and make the newly assembled dataset with MoRF annotations accessible to all developers: https://gsponerlab.msl.ubc.ca/software/morf_chibi/mc2.

## Author Contributions

**Nawar Malhis:** Dataset design and assembly; software design and implementation; writing (original draft, review, and editing). **Joerg Gsponer:** writing (review and editing); funding.

## Notes

https://github.com/NawarMalhis/MC2

### Competing Interest Statement

The authors have declared no competing interest.

### Summary of Updates

Updates to the test datasets and the writing.

